# Identification and characterization of a wet adhesive protein extracted from *Dreissena bugensis*, the freshwater quagga mussel

**DOI:** 10.1101/2024.10.16.618703

**Authors:** Angelico R. Obille, Rida Hasan, David J. Rees, Judith Ng, Karina M. M. Carneiro, Eli D. Sone

## Abstract

Mechanisms of wet adhesion have been developed by several aquatic organisms over millions of years of evolutionary processes. Yet, the repertoire of synthetic biocompatible wet adhesive materials is still limited. In most marine bioadhesive proteins, 3,4-dihydroxyphenylalanine (DOPA) plays a significant role in strong interfacial interactions. The bioadhesive proteins in freshwater organisms are less well understood. The quagga mussel (*Dreissena bugensis*) is a notorious freshwater invasive species in the Great Lakes that attaches to a plethora of surfaces via a byssus. To determine the adhesive proteins in the quagga mussel byssus, we utilized quantitative proteomics to identify the proteins enriched at the byssus-substrate interface. Among the identified proteins was the Dbfp7 protein family. Dbfp7 is a small, polymorphic, and mostly disordered protein that lacks significant amounts of DOPA. Atomic force microscopy measurements of Dbfp7 confirm that this protein has similar adhesion energy to marine mussel adhesive proteins in aqueous conditions despite lacking DOPA. These results suggest that freshwater mussels may employ different mechanisms of adhesion compared to marine byssates. The inclusion of Dbfp7 to the library of known wet bioadhesive proteins – the first functionally characterized freshwater bioadhesive protein to our knowledge – will allow for a better understanding of the fundamental properties required to achieve biocompatible wet adhesion, a crucial step for the development of bio-inspired wet adhesive materials, such as improved medical adhesives.

**Significance Statement:** While many aquatic organisms evolved strategies to adhere to surfaces underwater, current synthetic biocompatible adhesives lack reliable adhesive strength in varying aqueous conditions. Investigating freshwater mussel adhesive proteins expands the current understanding of aquatic bioadhesion and offers potential advancements in wet adhesive technology. The discovery of a DOPA-deficient wet adhesive protein derived from the invasive quagga mussels presents a paradigm shift to the currently known mechanisms of byssal bioadhesion, thereby expanding the repertoire of wet adhesive strategies. Further, understanding the mechanism of adhesion employed by this biofouling species can help with the design of anti-fouling strategies to mitigate the impacts of these animals on the infrastructure and ecosystems in the Great Lakes and connected freshwater systems in North America.

## Introduction

Adhesives found in nature perform in ways that synthetic adhesives have yet to match. For example, even though many aquatic organisms have adapted to adhere to surfaces underwater, it remains an engineering challenge to synthesize strong, biocompatible adhesives that are effective in wet conditions. One strategy that has been developed in nature over millions of years of natural selection is adherence via a structure called the byssus. The byssus is utilized by several bivalves, including marine mussels, oysters, and a few freshwater mussels, to permanently attach themselves and support their sessile lifestyle (1). The byssus is composed of individual proteinaceous threads extending from the body of the animal, with each thread ending distally in a flattened plaque. The bulk of the plaque material connects the thread to the adhesive proteins found at the plaque-substrate interface. These interfacial proteins bind so strongly that when plaques are detached, a residue called the footprint usually remains (2–5).

Originating from the Dnieper River in Eastern Europe and found more recently in the Great Lakes basin and connected waterways in North America, quagga mussels (*Dreissena bugensis*) and zebra mussels (*Dreissena polymorpha*) are invasive species that are among only a handful of freshwater byssates (6). Although superficially similar to that of the well-studied marine byssates, the biochemical composition of the *Dreissenid* byssus is unique; *Mytilid* byssal proteins and *Dreissenid* byssal proteins share very little homology (7, 8). Moreover, amounts of peptidyl 3,4- dihydroxyphenylalanine (DOPA) in *Dreissenid* byssal proteins are considerably lower than in marine mussels and are distributed evenly between the byssal thread and plaque rather than concentrated in the plaque (9). This suggests that in the context of protein-substrate interactions, *Dreissenids* may have evolved to employ a unique mechanism of adhesion that relies less heavily on DOPA-based interfacial interactions than their marine counterparts.

Byssal adhesive proteins that have been characterized in the current literature are predominantly in the marine context, i.e. in ocean environments with high ionic composition (1). The best studied system is the byssus of marine mussels of the genus *Mytilus* (10–14). Following the discovery of the significant role of DOPA in interfacial adhesion in marine mussels and other marine bioadhesive systems, DOPA-based adhesion has been established as a central paradigm for solutions to the problem of biocompatible wet adhesion (15–18). Subsequently, several materials utilizing DOPA- based chemistry for adhesion and cross-linking have been developed (19–24, 13, 25–29). Few of these materials, however, have translated to the clinical setting for medical adhesives, in part due to the propensity for catechol to oxidize into the non-adhesive DOPA-quinone form when exposed to pH values higher than 6.8 (30–32, 25). Aside from translational considerations, the direct effect of ionic strength in the adhesive mechanism of DOPA-based bioadhesion is still unclear (33–35). Studying bioadhesives from freshwater systems can help expand the current repertoire of biocompatible wet adhesives.

Sequencing of the quagga mussel foot transcriptome combined with bottom-up proteomics led to the determination of fourteen novel *Dreissena bugensis* foot protein sequences (Dbfp4 to Dbfp17), and more complete sequences of known Dbfp1 and Dbfp2 (8, 9, 36). The location and function of each protein, however, remains elusive. Of particular interest are the proteins at the plaque-substrate interface, where a 10 to 20 nm thick electron-dense footprint protein layer is present, as observed by transmission electron microscopy (TEM) (2). Here, we report the relative enrichment of the proteins in the bulk plaque and footprint via quantitative proteomics. This method allowed us to identify the proteins significantly enriched in the footprint, signifying potential adhesive ability. Among this set of footprint proteins is Dbfp7, a family of proteins that: matches the molecular weights of proteins detected by matrix-assisted laser desorption/ionization time-of-flight (MALDI-TOF) mass spectrometry (37), has been shown to be a significant portion of the byssus (8), and herein is shown to be adhesive in aqueous conditions despite lacking DOPA.

## Results

Quantitative liquid chromatography tandem mass spectrometry (LC-MS/MS) was used to identify the byssal proteins that are relatively more abundant in the footprint compared to the bulk plaque extracts. Quagga mussels collected from Lake Ontario were allowed to lay fresh byssi onto glass slides overnight. Footprint (FP) and bulk plaque (BP) proteins were separately solubilized in extraction buffer following scraping of the bulk plaque, which leaves behind a “footprint” residue on the slide (**Figure 1A**) (3–5, 8, 9). The footprint and bulk plaque solubilized proteins were digested with trypsin and the peptides were subsequently functionalized with tandem mass tag (TMT) labels. The tagged peptides were pooled then sequenced via LC-MS/MS (**Figure 1B**). The peptide sequences were fingerprinted against the quagga mussel foot transcriptome (8), searched against protein databases using the basic local alignment search tool (BLAST) algorithm for homology assessment and identification of proteins, and compared to proteins identified from whole-thread/plaque byssal material that was induced to be secreted by potassium chloride (KCl) injection in the pedal ganglion (IND); the induced byssus dataset has been previously reported (8).

**Figure 1:**
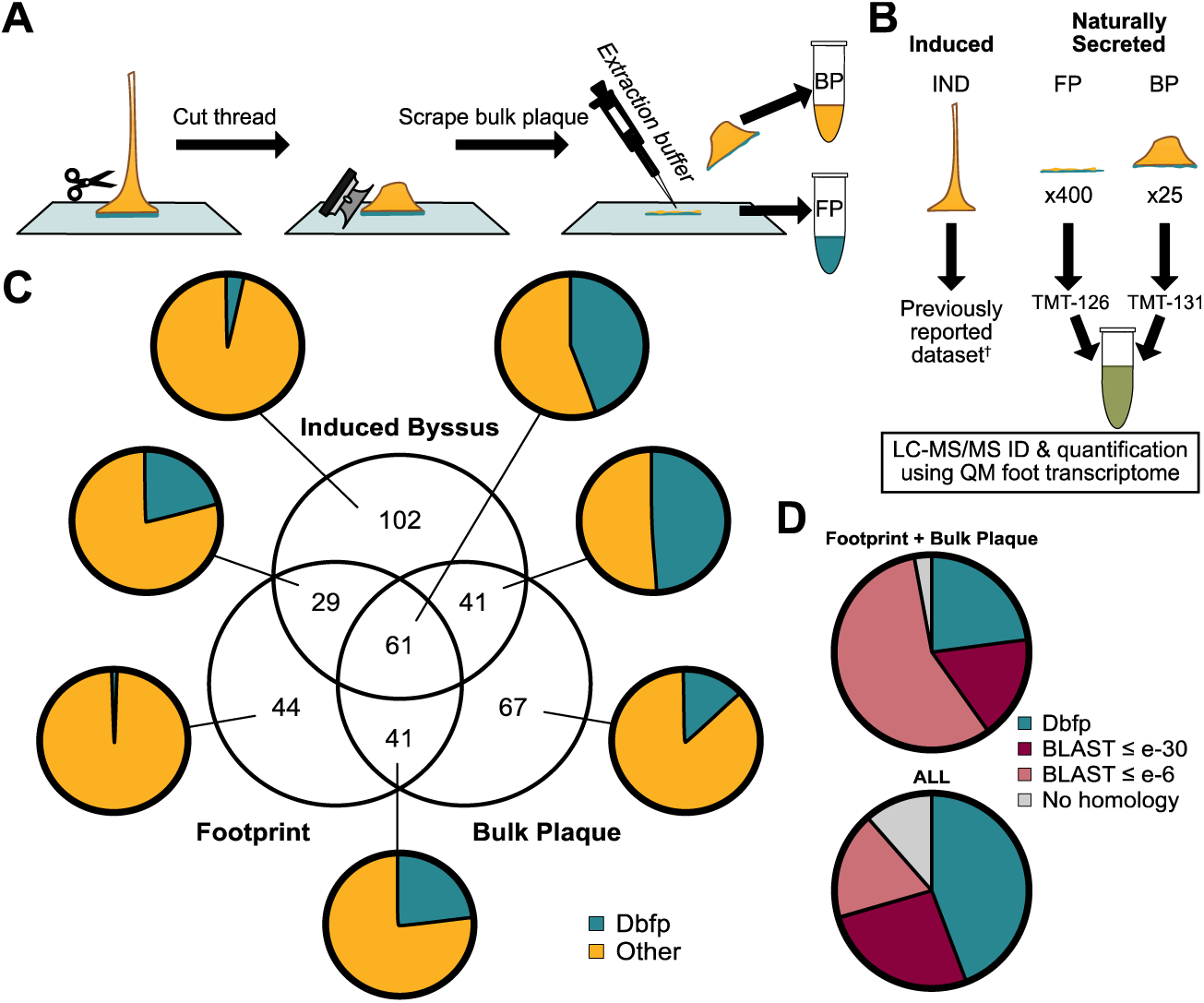
Overview of plaque proteomics investigation. (A) Schematic of sample collection. (B) Summary of workflow. (C) Number of unique proteins found in each group of extractions. Insets indicate the proportion of Dbfp’s in each group. (D) Distribution of proteins in the intersecting ‘footprint + bulk plaque’ group and the ‘ALL’ group, highlighting the proportion of proteins that are Dbfp’s or show strong homology (BLAST score ≤ e^-30^), low homology (BLAST score ≤ e^-6^), or no homology to known proteins. †(8)

### Identification of plaque proteins in the naturally secreted quagga mussel byssus

The induced byssus includes proteins composing the entire byssus, i.e. thread proteins, bulk plaque proteins, and footprint proteins. In the naturally secreted samples, bulk plaque and footprint samples were collected in separate groups, and the threads were discarded. The plaque protein collection method cannot precisely separate the bulk plaque from the footprint material. As a result, footprint proteins are expected to be present in the bulk plaque group (albeit to a lesser extent) and vice versa. For this reason, the ‘Footprint + Bulk Plaque’ group and the ‘ALL’ group were considered to most reliably contain proteins localized to the plaque (**Figure 1D**). These proteins were identified by the BLASTp algorithm and include (full list in **Table S1**): Dbfp’s and proteins with high homology to known enzymes, structural proteins, matrix proteins, protease inhibitors, mucin-related proteins, immune/defense proteins, calcium-binding protein, and rhamnose-binding protein.

### Quantitative proteomics classifies 21 footprint-enriched byssal proteins

Several proteins were found to be enriched in the footprint relative to the bulk plaque (**Figure 2**). The proteins that were measured to have a footprint-to-bulk plaque fold change ratio (FCR footprint : bulk plaque) > 2 that is statistically significant (p-value < 0.05) are likely to play direct roles in the adhesive mechanism since they are enriched at the interface between the byssus and the substrate.

**Figure 2:**
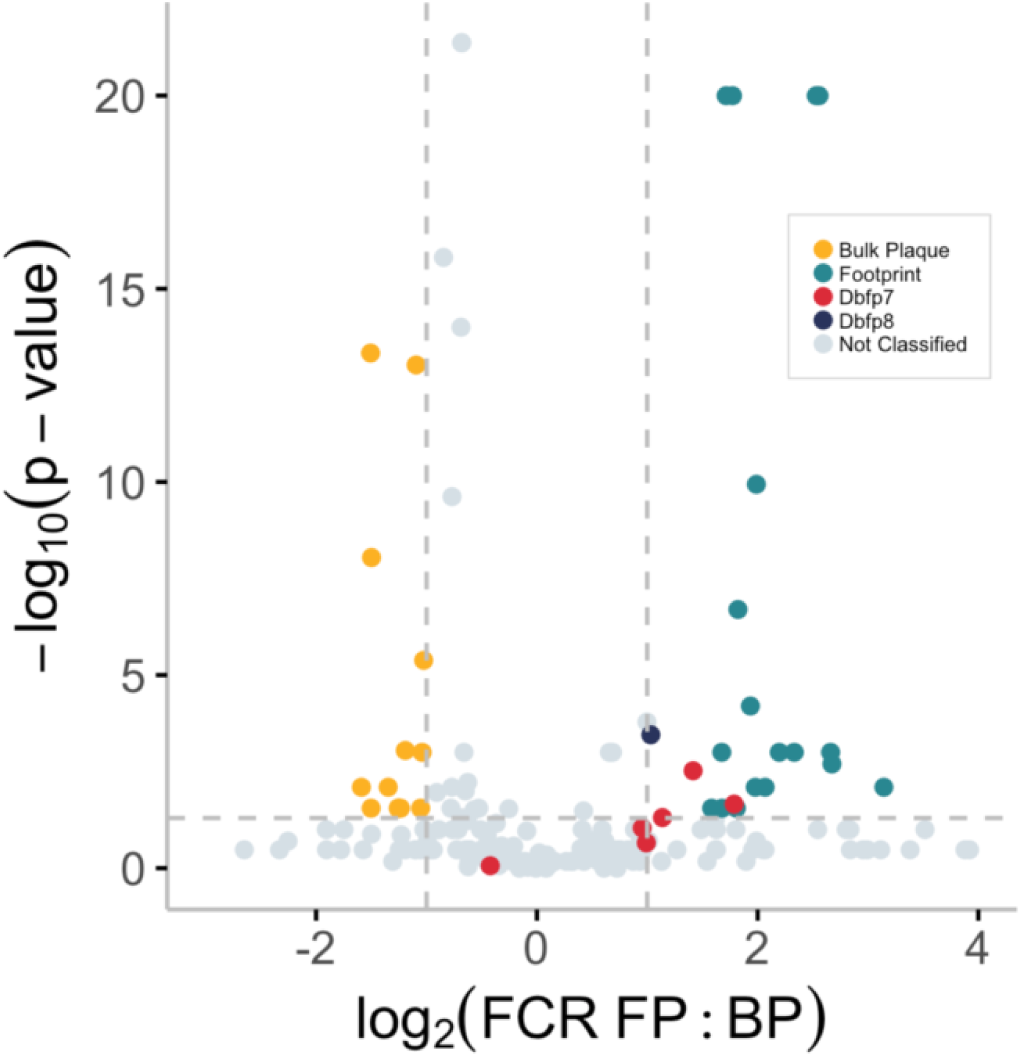
Volcano plot of quagga mussel plaque proteins. Each point represents an identified protein. Proteins with a log2(fold change ratio between footprint and bulk plaque) > 1 (i.e. > 2 FCR) and a −log10(p-value) > 1.3 (i.e. p-value < 0.05) are significantly enriched in the footprint. Dbfp7 variants and Dbfp8 are indicated as the red and blue points, respectively.

There were 22 significantly enriched footprint proteins identified (**Table 1**). Among these are three variants of Dbfp7 and one variant of Dbfp8. The other proteins are weakly (e-score > e-6), moderately (e-30 < e-score < e-6), or strongly (e-score < e-30) homologous to known proteins, identified by utilizing the BLASTp algorithm against a clustered non-redundant database (38, 39). Overall, the proteins relatively enriched at the adhesive interface are: Dbfp7, Dbfp8, kielin/chordin-like protein, zonadhesin/IgGFc- binding protein, neurotrypsin-like protein, mucin-like protein, peroxidase, plasminogen-like protein, galaxin-like protein, and several homologues to uncharacterized proteins from other aquatic organisms. Interestingly, some of the proteins contained at least one possible hydroxylated tyrosine residue (i.e. DOPA). None of these proteins, however, were notably abundant with DOPA, corroborating the overall scarcity of DOPA in *Dreissenid* byssus (9) or the limited ability to solubilize DOPA-containing proteins, which can cross-link post-secretion (16, 40–43). Dbfp1 and Dbfp2, previously identified to have the most DOPA (9), were not present in the footprint nor bulk plaque. One transcript (comp50166_c1_seq2-2R) did not have a clear protein-coding open reading frame and was thus excluded as a footprint protein. Parts of the identified peptides of this transcript are probably components of a different protein with a similar transcript sequence.

**Table 1:** The quagga mussel footprint proteome in order of relative enrichment in the footprint. Accession #: transcript label and reading frame (0F, 1F, 2F, 0R, 1R, or 2R). log2(FCR FP:BP): log-transformed fold change ratio between footprint and bulk plaque. −log10(p-value): log-transformed significance value. SP: signal peptide. DOPA: confident (Y), possible (M), or no (N) detection of at least one hydroxylated tyrosine (peptidyl 3,4-dihydroxyphenylalanine) residue via LC-MS/MS sequencing. pI: isoelectric point calculated from the translated transcript sequence. MW: molecular weight calculated from the translated transcript sequence.

| *Identification* | *Accession #* | *$\log_2(\text{FCR FP:BP})$* | *$-\log_{10}(\text{p-value})$* | *SP? (Y/N)* | *DOPA? (Y/M/N)* | *pl* | *MW (kDa)* |
| --- | --- | --- | --- | --- | --- | --- | --- |
| Kielin/Chordin-like | comp50443_c0_seq4-0F | 3.14 | 2.10 | Y | M | 7.99 | 15.2 |
| Mucin/Spondin-like | comp52859_c0_seq1-1R | 2.67 | 2.70 | N | N | 4.59 | 113 |
| Hornerin | comp41116_c0_seq1-2F | 2.66 | 3.00 | Y | N | 8.06 | 8.26 |
| Zonadhesin/IgGFC-binding | comp53519_c2_seq1-2R | 2.56 | 20.0 | N | M | 5.31 | 125 |
| Zonadhesin/IgGFC-binding | comp53519_c2_seq2-0R | 2.53 | 20.0 | N | N | 5.25 | 128 |
| Hornerin | comp36226_c0_seq1-1F | 2.33 | 3.00 | Y | N | 8.48 | 16.3 |
| CD109 antigen-like | comp30100_c0_seq2-2F | 2.20 | 3.00 | Y | N | 4.88 | 28.2 |
| TPF1III | comp56420_c11_seq1-2F | 2.07 | 2.10 | N | N | 8.01 | 8.67 |
| Kielin/Chordin-like | comp31275_c0_seq2-2R | 1.99 | 9.94 | N | M | 8.16 | 5.51 |
| TTN1 | comp53519_c2_seq3-0R | 1.98 | 2.10 | N | N | 4.76 | 35.7 |
| Peroxidase | comp51774_c0_seq1-2F | 1.94 | 4.20 | N | N | 8.37 | 83.0 |
| Kielin/Chordin-like | comp31275_c0_seq1-2R | 1.82 | 6.70 | Y | M | 5.18 | 13.9 |
| Kielin/Chordin-like | comp30932_c0_seq1-0F | 1.81 | 1.55 | N | N | 7.53 | 9.08 |
| Dbfp7 | comp52765_c1_seq12-1F | 1.79 | 1.66 | Y | Y | 9.22 | 12.0 |
| Fibrillin/Galaxin | comp56544_c0_seq1-2R | 1.77 | 20.0 | N | M | 5.95 | 149 |
| Fibrillin/Galaxin | comp56544_c3_seq1-2R | 1.71 | 20.0 | N | M | 6.91 | 231 |
| Unknown | comp57807_c0_seq1-0R | 1.68 | 1.55 | N | N | 8.82 | 12.4 |
| Galaxin | comp56544_c8_seq1-1R | 1.67 | 3.00 | N | N | 5.84 | 29.7 |
| Dbfp7 | comp52765_c1_seq11-1F | 1.42 | 2.52 | Y | Y | 8.83 | 9.44 |
| Dbfp7 | comp52765_c1_seq14-1F | 1.14 | 1.31 | Y | Y | 8.80 | 9.90 |
| Dbfp8 | comp44862_c1_seq1-2F | 1.03 | 3.45 | Y | M | 9.99 | 16.4 |

### Selection of Dbfp7 as the first footprint protein to characterize

Previously, three quagga mussel byssal proteins were identified via a DOPA-specific stain on protein extracted from quagga mussel feet and named Dbfp1, Dbfp2, and Dbfp3 (9). Subsequent characterization of Dbfp1, one of the higher molecular weight DOPA-proteins, has classified it as a cuticle protein or a structural cohesive component (9, 44, 36). The role of Dbfp2 remains unclear (8, 9, 45) and no investigation of Dbfp3 has been done to date.

Proteins of similar molecular weight as Dbfp3 were detected in the plaque by MALDI-TOF mass spectrometry and were inferred to contain little to no DOPA (37). Dbfp3 was reclassified as Dbfp7 upon the construction of the quagga mussel foot transcriptome, which revealed significant homology of this protein to the known zebra mussel protein Dpfp7 (8, 46). Herein, the Dbfp7 protein family was identified to be among the proteins that are significantly enriched in the footprint. Taken together this suggests that Dbfp7 plays an important role in byssal adhesion, potentially having a role in bonding to a surface. Dbfp7 was thus selected as the first adhesive protein candidate to be extracted and further characterized.

### Dbfp7 is polymorphic and DOPA-deficient

Dbfp7 was extracted from quagga mussel phenol glands (∼ 5 µg per gland) via several steps of selective precipitation and reversed-phase high-pressure liquid chromatography (HPLC) (**Figure S1**) (9). The most abundant residue of Dbfp7 is glycine, followed by significant amounts of proline, asparagine, and tyrosine. There are 13 known variants of Dbfp7 detected in naturally secreted byssi, induced byssi, and purified Dbfp7, with lengths ranging from 82 to 132 residues (**Figure 3A**). These variants can be broadly classified as either acidic or basic variants, determined by the theoretical isoelectric point of the sequence. The acidic variants all contain a 28-residue sequence in the C-terminal half of the protein sequence, which is missing from the basic variants. Most of the variation of the Dbfp7 sequence occurs within the first 50 residues; the C-terminal half of Dbfp7 is highly conserved (aside from the presence or absence of the 28-residue sequence).

**Figure 3:**
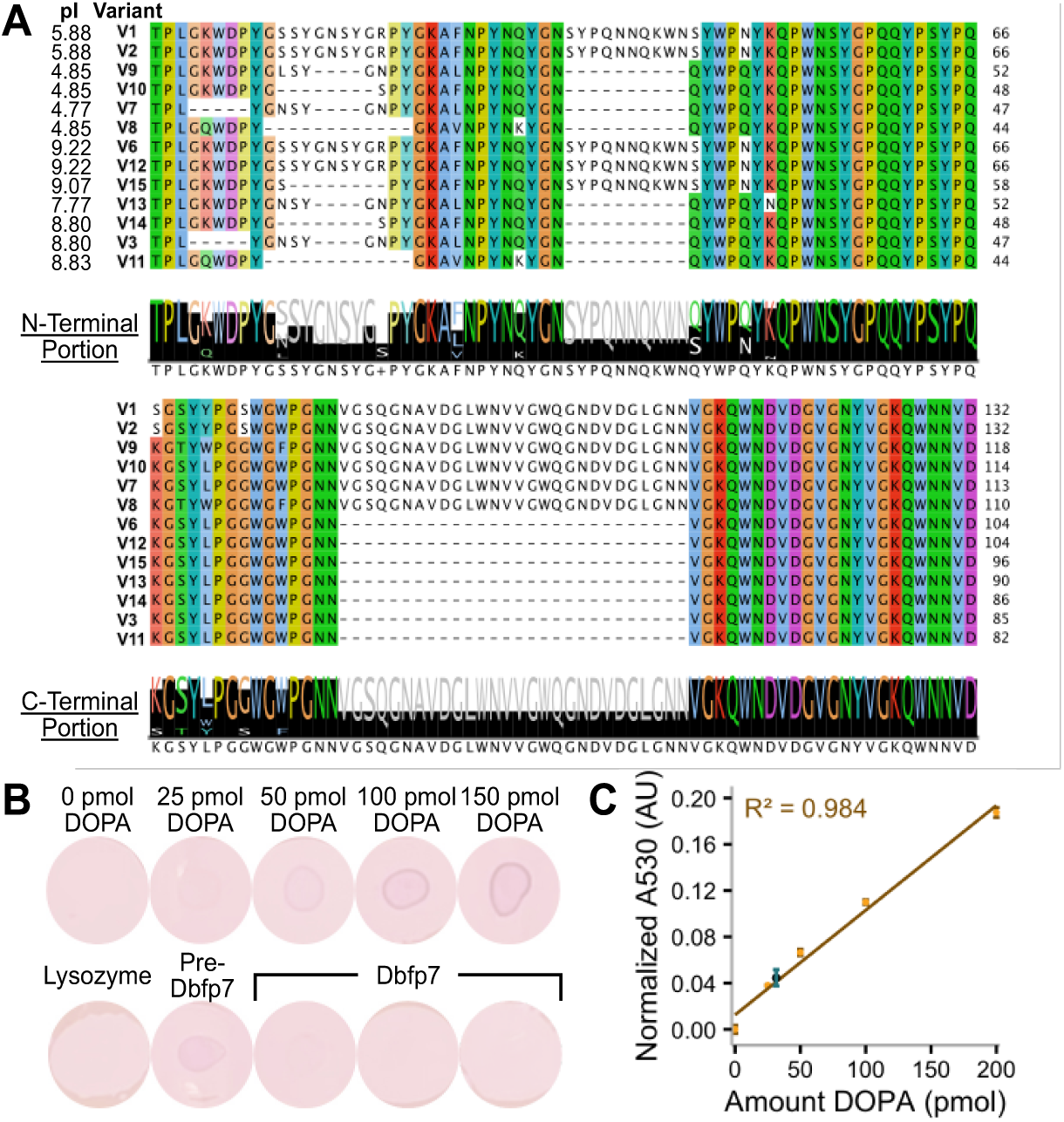
The Dbfp7 protein family is rich in glycine, proline, asparagine, and tyrosine and lacks significant amounts of DOPA. (A) Sequences of Dbfp7 variants. Variants are sorted by length, demonstrating the 28-residue region that is present only in the C-terminal half of the acidic variants. The consensus chart (generated by Jalview 2.11.3.3) below the alignment indicates the residues that are highly conserved, particularly in the C-terminal portion. (B) Dot blot with known amounts of peptidyl DOPA, lysozyme negative control, pre-Dbfp7 HPLC fraction, and Dbfp7 stained with NBT. (C) Calibration curve obtained with known amounts of peptidyl DOPA; the blue point indicates the absorbance of Dbfp7, corresponding to 0.11 ± 0.02 mol% DOPA. Error bars indicate standard deviation (n = 3).

The amount of DOPA in Dbfp7 was found to be notably scarce. Only 1-2 LC-MS/MS sequences of Dbfp7 peptides (in either purified, naturally secreted, or induced instances) confidently contained DOPA, depending on the variant and the extracted pool. The same site of tyrosine hydroxylation was not conserved between induced, purified (pre-secreted), and naturally secreted forms. Most detected hydroxylation modifications were ambiguous (**Figure S4**). Specifically, the identified putative DOPA residues were in spectra of peptides that were not fully fragmented; thus, it is possible the detected hydroxylation was on a different residue within the peptide (e.g. hydroxylysine). Further, purified Dbfp7 is only slightly positively stained with nitroblue tetrazolium (NBT) (**Figure 3B**). The amount of DOPA in Dbfp7 was calculated to be 0.11 ± 0.02 mol% (n = 3, s.d.), about 1 DOPA residue per 10 Dbfp7 molecules (**Figure 3C**). It is possible that Dbfp7 and other plaque proteins are hydroxylated extracellularly by tyrosinase, which has been found in the bulk plaque (**Table S1**). Interestingly, the chromatographic fraction of the S2 mixture (**Figure S5C**) eluted before the Dbfp7 elution (40-45% acetonitrile), i.e. “pre-Dbfp7 HPLC fraction”, is NBT-positive (**Figure 3B** and **Figure S5A**). This signifies that DOPA can be detected following the extraction method via HPLC. Gel electrophoretic separation of this fraction reveals the presence of low molecular weight proteins with similar weights of purified Dbfp7 (**Figure S5B**). Thus, it is possible that the proteins in the strong low molecular weight NBT-positive band detected by Rzepecki and Waite (9), so-called Dbfp3, was not primarily Dbfp7, but rather was mostly a different protein in the extracted mixture. In any case, the population of Dbfp7 extracted from phenol glands (pre-secretion) characterized herein is deficient in DOPA.

Purified forms of Dbfp7 from phenol glands are significantly lower in molecular weight than that of the full theoretical length of the sequence despite the detection of peptides from the full sequence within the purified sample (**Figure S1** and **Figure S2**). Curiously, the localization method of Dbfp7 reveals that although it is overall relatively more abundant in the footprint, the peptides of Dbfp7 detected in the bulk plaque consistently map to the C-terminal portion and the peptides in the footprint mostly map to the N-terminal portion (**Figure S3**). This suggests that Dbfp7 is post-translationally or post-secretionally cleaved. The population of Dbfp7 analyzed herein, which is extracted pre-secretion, also does not contain the full-length molecule (**Figure S2**), therefore, Dbfp7 is likely to be cleaved intracellularly. This raises the possibility that the two portions of Dbfp7 have differing functions, where the N-terminal region interacts with substrates while the C-terminal region is responsible for protein-protein interactions resulting in cohesion between the adhesive and the bulk plaque material. Alternatively, it is possible that C-terminal peptides interacting with the substrate were unable to be extracted from the footprint.

Preliminary structural investigations of Dbfp7 reveal both ordered and disordered structure. Due to the presence of multiple variants, structural measurements are an average of the Dbfp7 population extracted from phenol glands. To characterize the structure of Dbfp7, circular dichroism (CD) spectra of Dbfp7 dissolved in 5% acetic acid (**Figure 4A**) were deconvoluted by Contin-LL (47). The proportions of structural features were 3.3 ± 0.3% ɑ-helix, 33 ± 5% β-strand, 20 ± 3% β-turns, and 44 ± 5% disordered (**Figure 4B**). While almost half of the protein was found to be disordered, significant portions of the sample adopted β-strand and β-turn structures. To predict the structure, the AlphaFold protein folding algorithm was used. Due to caveats imposed by the lack of homology of Dbfp7 to other known proteins, pLDDT scores used to determine model confidence were very low (<0.5). The structure is predicted to be entirely disordered; however, the low homology of Dbfp7 to known proteins may result in the prediction algorithm missing ordered regions. Qualitative observations of the model show a C-terminal region that is slightly more compact than the N-terminal portion, which looks relatively extended (**Figure 4C**).

**Figure 4:**
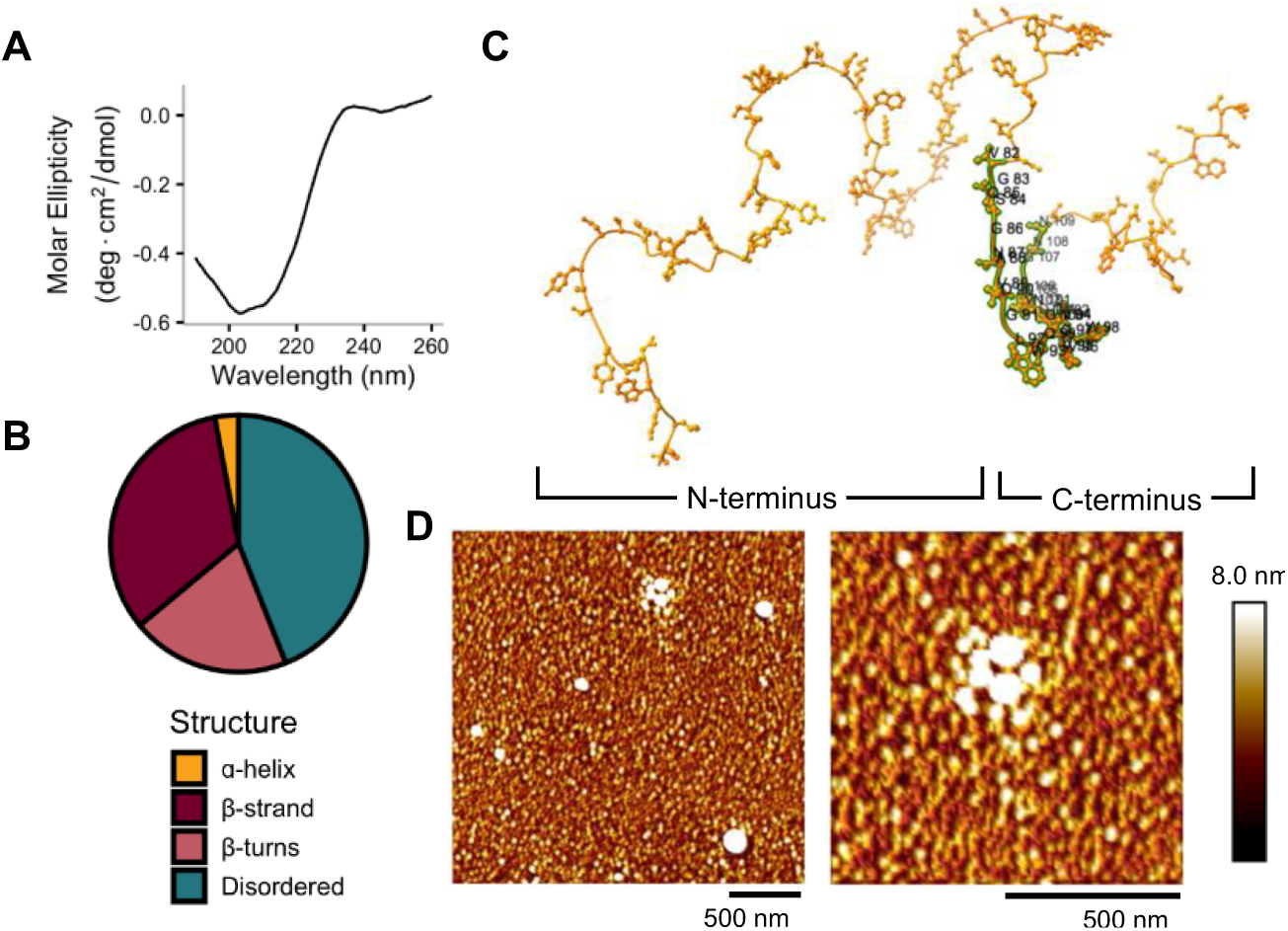
Dbfp7 surface morphology and structure. (A) CD spectrum of Dbfp7 dissolved in 5% acetic acid. (B) Proportion of secondary structure features via deconvolution of CD spectrum by Contin-LL. (C) Predicted structure of Dbfp7 (variant 1) visualized by ChimeraX. Labelled residues indicate the 28-residue C-terminal region found only in acidic variants. (D) Atomic force microscopy height images of Dbfp7 on mica in aqueous conditions. Scalebars each represent 500 nm.

### Dbfp7 is adhesive in aqueous conditions

To assess morphology on a surface and adhesive ability, Dbfp7 was drop casted onto freshly cleaved mica and imaged in aqueous conditions using an atomic force microscope (AFM) in PeakForce tapping mode with quantitative nanomechanical mapping (PFQNM). At 100 µg/mL, a dense layer of round and flat globules (ranging from 3.5 to 35 nm in height) was found throughout the surface (**Figure 4D**).

The surface containing Dbfp7 was adhesive, with some portions of relatively lower measured adhesion force that correspond to larger globules in the height map (**Figure 4D** and **Figure 5A**). Overall, Dbfp7 is relatively more adhesive than the control protein, bovine serum albumin (BSA), known to be non-adhesive (**Figure 5C**) (48–51). The average forces of adhesion for Dbfp7 and BSA were measured to be 470 ± 160 pN and 160 ± 140 pN, respectively (s.d. and n = 262,144). Interestingly, the space between Dbfp7 globules (so-called interglobular space) is more adhesive than the globules themselves (**Figure 5B**). This may be due to the inconsistent indentation pressure during scanning, since the setpoint was kept constant during data acquisition.

**Figure 5:**
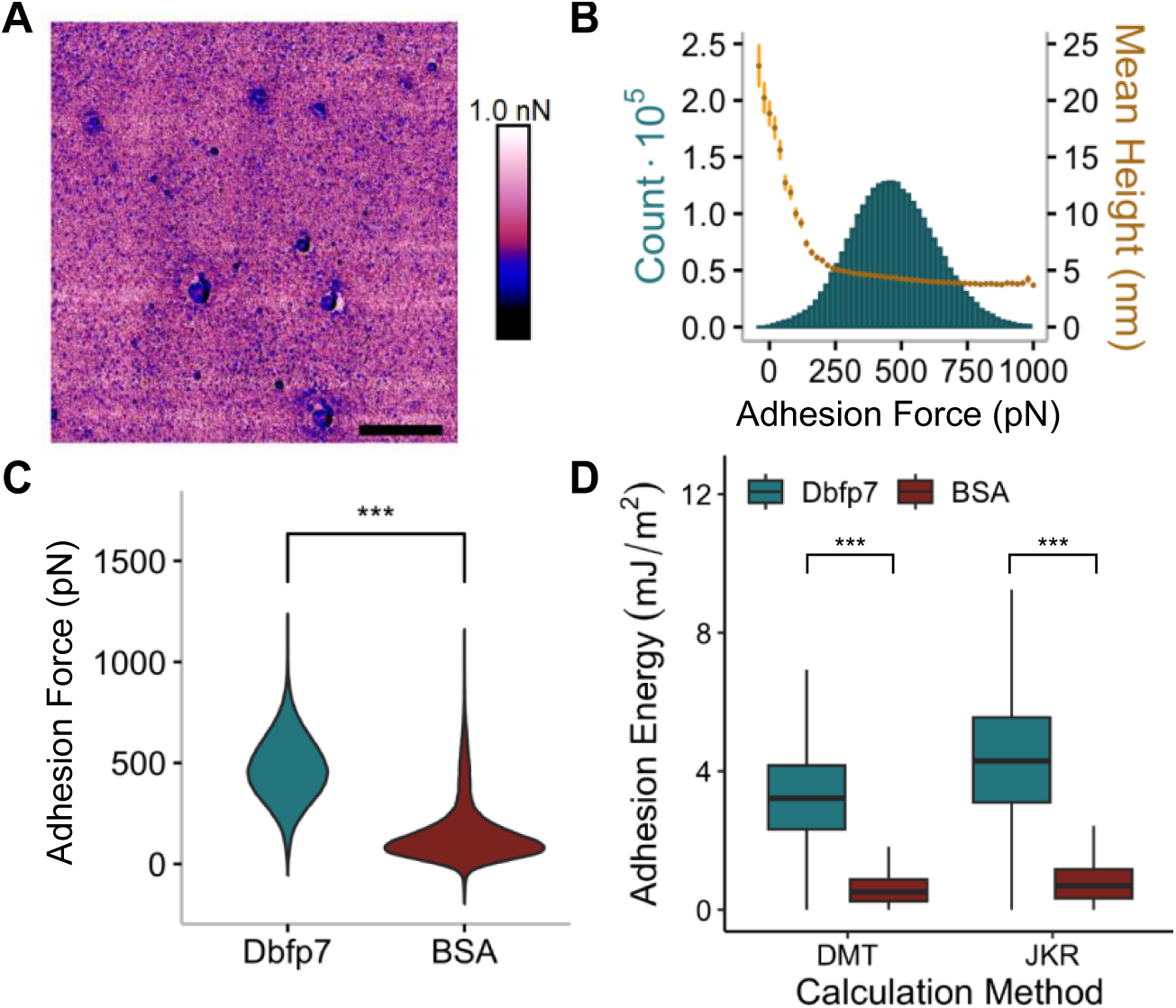
Dbfp7 is adhesive in an aqueous environment. (A) Adhesion map of Dbfp7 (100 µg/mL) coated on mica, imaged in aqueous conditions. Scale bar is 1 µm. (B) The mean height of each bin in the adhesion histogram demonstrates that the areas of low adhesion correspond to regions with taller features. Error bars indicate the standard error of the mean (s.e.m.). (C) Dbfp7 is overall more adhesive than samples prepared with BSA (n = 262,144). (D) Adhesion energy of Dbfp7 and BSA calculated by the DMT and JKR contact models from adhesion force measurements obtained by PFQNM AFM (s.d., N = 2,859,269 and 793,084 for Dbfp7 and BSA, respectively). *** indicates p-value < 0.001.

Alternatively, the interglobular space may contain single molecules of Dbfp7 that are more readily available for adhesion to the AFM tip compared to self-assembled globules. This trend is not present in BSA samples (**Figure S6**).

The DMT (Derjaguin-Muller-Toporov) and JKR (Johnson-Kendall-Roberts) models of adhesion predict different contact areas between two surfaces depending on the deformation of the indenting probe (52–54). Depending on the model used, the adhesion energy from the measured adhesion force may vary, and therefore, the appropriate model is required to convert the measured adhesion force to adhesion energy (55–57). The Tabor parameter approximates the appropriate model based on the radius and moduli of the surfaces allowing for the determination of the appropriate theory for the materials in the analyzed system. The Tabor parameter for this experimental set-up was calculated to be ∼0.12 (see Supplemental Information for full calculation details). This value means that the contact in this system lies between the characteristic contact regimes described by the two models. Currently, there is no model to adequately describe a system that is characteristic of contact partially described by either the DMT or JKR contact models. Therefore, we report the converted adhesion energy using both the DMT and the JKR models (**Figure 5D**) to compare to adhesive proteins in current literature. The calculated adhesion energy of Dbfp7 is 4.4 ± 1.8 mJ/m2 (JKR) or 3.3 ± 1.3 mJ/m2 (DMT) which is ∼5 times higher than BSA (0.90 ± 0.89 mJ/m2 (JKR) or 0.67 ± 0.66 mJ/m2 (DMT)) (s.d., N = 2,859,269 and 793,084 for Dbfp7 and BSA, respectively). Interestingly, the adhesion energy value for Dbfp7 is in a similar range to the adhesion energy of Mfp3, one of the *Mytilus* adhesive proteins (∼3-6 mJ/m2 determined by surface forces apparatus (SFA) measurements) (14). However, the other known *Mytilus* interfacial protein, Mfp5, remains the strongest known byssal adhesive protein with the most relative amount of DOPA (15 mJ/m2 and 30 mol% DOPA) (14).

## Discussion

Several plaque proteins were identified to be relatively more enriched in the footprint than in the bulk plaque material, suggesting they may play direct roles in plaque-substrate adhesion. Among these, Dbfp7 stood out due to its detection in previous studies on quagga mussel byssal proteins. When extracted from phenol glands (pre-secretion), Dbfp7 adopts a disordered structure with significant proportions of β- strand and β-turn features and contains little to no DOPA. Despite this lack of DOPA, adhesion measurements by AFM performed in aqueous conditions revealed that Dbfp7 exhibits wet adhesion similar to Mfp3, one of the marine mussel adhesive proteins.

Dbfp7 is thus classified as a DOPA-deficient wet adhesive protein.

The marine *Mytilids* and the freshwater *Dreissenids* evolutionarily diverged at the class level; *Dreissenids* have since evolved in brackish water originally and eventually adapted to freshwater habitats (58). It is evident that the significant amount of time between common ancestors gives rise to a variety of differences in the byssal adhesion mechanism. For example, another freshwater byssate *Limnoperna fortunei* (golden mussel) – found in the freshwater systems of Japan, Korea, and Eastern China – diverges from *Mytilus sp*. at the family level and is thus genetically more similar. Indeed, it has been found that the golden mussel byssus contains typical marine mussel adhesive protein domains such as EGF domains, vWFA domains, and GXX motifs (59). Further, DOPA was found in the outer sheath of golden mussel byssal threads and at the bottom of adhesive plaques by NBT staining (59). The presence of DOPA at the adhesive interface suggests DOPA-based interfacial chemistry, however the specific golden mussel footprint proteins have not been characterized yet. In the zebra mussel byssus, DOPA and catechol oxidase was found throughout the plaque, suggesting that some interfacial *Dreissenid* footprint proteins may employ DOPA interfacial chemistry while bulk plaque and thread DOPA-proteins contribute to cross-linking (60). However, the adhesion mechanism of Dbfp7 may be less dependent on DOPA, given the even distribution of DOPA between the thread and plaque.

In the marine mussel, the proteins found at the plaque-substrate interface have significantly higher relative amounts of DOPA compared to the bulk plaque and thread proteins. In *Dreissenids*, it was found that peptidyl-DOPA is present in low amounts throughout the byssus, including the thread (9). This suggests that the role of DOPA in the freshwater system is primarily for cross-linking purposes (16, 40–43). Dbfp7 purified from phenol glands of quagga mussels has herein been shown to lack significant amounts of DOPA while being more adhesive than BSA under the same conditions (**Figure 5**). Conversion of the adhesive forces to adhesion energy reveals similar adhesion energy of Dbfp7 compared to one of the marine mussel interfacial proteins, Mfp3. Although similar in molecular weight with abundant glycine, tyrosine, proline, and asparagine residue proportions, Dbfp7 and Mfp3 share very little sequence homology and Mfp3 is more enriched with DOPA (around 10-20 mol%) compared to Dbfp7 (∼0.1 mol%). The other known interfacial *Mytilus* protein, Mfp5, contains even more DOPA (30 mol%) than Mfp3 and is significantly more adhesive than both Mfp3 and Dbfp7 (15 mJ/m2 measured by SFA) (14). It appears that some tyrosine residues in Dbfp7 may be post-secretionally hydroxylated within the distal depression and secreted byssus, since tyrosinases were identified in the bulk plaque (**Table S1**). Still, the DOPA-deficient pre-secretion population of Dbfp7 is adhesive.

The mechanism of adhesion for Dbfp7 remains unknown and further studies into specific peptide sequences and post-translational modifications are required to develop a model for the adhesion mechanism. It is possible that in environments with lower ionic strength, as in freshwater habitats, lower amounts of DOPA can achieve similar adhesive strength. Further, several efforts are being made to understand the role of lysine and other cationic moieties in conjunction with DOPA-based adhesion, particularly in cation-pi interactions for cross-linking and complex coacervation (34, 61–67). It is unclear whether the dihydroxyl moiety in catechol is necessary for this interaction or if the phenyl moiety in residues such as tyrosine, phenylalanine, and tryptophan is sufficient (68, 69). Charged residues may also play a role in dislodging surface-bound water molecules and surface ions to allow for direct contact with a surface (34). Indeed, Dbfp7 contains significant amounts of aromatic residues tyrosine and tryptophan, as well as positively charged lysine (**Figure 3A**).

Dbfp7 is polymorphic with up to 13 detected variants. Several genes of the zebra mussel homologue of Dbfp7, Dpfp7, have been identified in multiple regions of the zebra mussel genome (70). Mfp3 is similarly polymorphic, with up to 35 known variants encoded throughout the *Mytilus sp.* genome (5). The existence of multiple variants of a gene that is preserved several times in the genome suggests vital importance.

Particularly, polymorphism of an interfacial protein may enhance the versatility of adhesion to different kinds of surfaces (3, 5, 8, 16). Further, Mfp3 is known to exist as two populations, Mfp3S and Mfp3F, so-called the slow and fast variants due to their elution time in reversed-phase HPLC, thereby differing in hydrophobicity. Mfp3S was found to be more resistant to oxidation and retains adhesion when pH is equilibrated to seawater levels (7, 71). The role of the two variant types of Dbfp7 (acidic and basic) is currently unclear and may: (i) function similarly to the Mfp3 variants, (ii) have different roles in the footprint-bulk plaque and footprint-substrate interfaces, or (iii) contribute to spreading behaviour on a surface by the formation of complex coacervates during secretion (72–74). Encoding and synthesizing these two peptide components in the same transcript may have the advantage of maintaining stoichiometric proportions.

The quagga mussel footprint contains 20 proteins in addition to Dbfp7 that contribute to byssal adhesion, possibly in interfacial interactions (**Table 1**). Further investigation is needed to understand the interactions between these proteins with each other, with bulk plaque proteins, and with adherend substrates. Of particular interest are the zonadhesin/IgGFc-binding proteins, homologues of which were found in the sea star (*Asteria rubens*) adhesive footprint (75). Further, kielin/chordin-like proteins match the molecular weights of interfacial plaque proteins identified by MALDI-TOF (37). With the advent of high-throughput sequencing techniques, the library of proteins found in bioadhesive systems is vast. Little has been done to localize these proteins at the direct adhesive-substrate interface *in situ*, with a larger current focus on bulk cohesive properties than on interfacial adhesion chemistry. Localization of proteins at direct adhesive-substrate interfaces will provide a smaller subset to analyze for understanding the fundamental mechanisms of bioadhesion in aqueous environments.

It is difficult to directly compare adhesive measurements of Dbfp7 to other studies since the methods of wet adhesive protein characterization are varied. SFA developed by Israelachvili and applied by Waite and others to marine byssal proteins has been the predominant method of adhesive measurements (14, 34, 69, 76–79). This unique instrument is not as widely implemented compared to AFM. Although similar in fundamental operation (i.e. generation and analysis of submicron contact force-distance curves), the planar scale of these two systems is different, with SFA analyzing at the squared milli- to centimetre contact area range and AFM analyzing at the squared nano- to micrometre contact area range. The classical models of contact (Hertz, DMT, JKR, etc.) may allow for the comparison of adhesion energies obtained from different systems; however, the physics at these two scales may be fundamentally different and thus may not be directly comparable. To date, the adhesion in systems that lie between DMT and JKR contact models is yet to be fully understood (55, 56, 80, 81). Further complications arise when the two contacting surfaces are not entirely flat or smooth (57, 82). Advancements in theoretical contact physics at the nanoscale are needed to harmonize any discrepancies that may arise from measuring adhesion at differing size scales. The problem of scale may be circumvented with the use of colloidal probe technique, allowing for the analysis of adhesion at the micrometre contact area scale in AFM (83, 84). This technique, however, cannot be used for concurrent imaging and characterization of mechanical properties. Further, the adhesion measurements obtained herein presumably represent adhesion of Dbfp7 with the silicon nitride AFM tip. It is possible that some adhesion measurements were from Dbfp7 bound to the tip, resulting in a measurement of the interactions between Dbfp7-to-Dbfp7 or Dbfp7-to- mica. Since the protein seems to aggregate resulting in a non-uniform molecular layer and due to the limited lateral resolution of AFM imaging, these possible interactions may contribute to the observed adhesion average.

This study highlights the importance of studying aqueous bioadhesive proteins from a diversity of organisms. To our knowledge, Dbfp7 is the first freshwater wet adhesive protein to be functionally characterized. The lack of DOPA in this protein is interesting but not shocking when considering the evolutionary history of *Dreissenids* compared to the other bioadhesive systems studied so far. The inclusion of Dbfp7 (and potentially others from the 20 footprint proteins herein localized) to the repertoire of wet bioadhesive proteins will allow for the identification of the fundamental properties required to achieve biocompatible wet adhesion for the development of bio-inspired wet adhesive materials. Further, understanding the specific mechanism of adhesion employed by *Dreissenids* that are invasive species to the Great Lakes basin and connected waterways in North America will aid in the development of anti-fouling strategies (85).

## Materials and Methods

### Quagga mussel collection and maintenance

Freshwater mussels were harvested from Lake Ontario at Port Credit Harbour Marina in Mississauga, Ontario. Live mussels were cleaned and sorted by species (quagga mussels and zebra mussels) and transferred to tanks with circulating artificial lake water (0.30 mM sodium bicarbonate, 0.13 mM potassium bicarbonate, 0.35 mM sodium chloride, 0.67 mM magnesium sulfate, 0.08 mM magnesium chloride, 1.47 mM calcium carbonate) at 12°C and fed powdered green algae.

### Plaque protein collection

Quagga mussels were first detached from their original substrates (tank walls or other mussels), secured into individual cages with glass slides, and allowed to lay byssi overnight. Mussels were gently detached from the substrates by cutting the byssal threads, leaving the plaque intact on the glass slide. Bulk plaque material was scraped off with a razor blade and collected in a pool of extraction buffer (composed of 200 mM sodium borate, 4 M urea, 1 mM potassium cyanide, 1 mM ethylenediaminetetraacetic acid, and 10 mM ascorbic acid, at pH 8.0). Extraction buffer was deposited onto the footprint residue remaining on the slide, incubated for 1-2 minutes, then collected.

Samples were homogenized in a 1 mL tissue grinder on ice, centrifuged at 17,000 x g for eight minutes at 4°C, then flash frozen for storage until analysis. Amounts of bulk plaque and footprint pools were normalized by weight (resulting in 25 bulk plaques and 400 footprints in total for comparative analysis).

### Tandem mass spectrometry on digested peptides extracted from freshly secreted byssus

The bulk plaque and the footprint pools were digested with 13 ng/µL trypsin (Porcine, sequencing grade, Promega) overnight at 37°C. Following digestion, the bulk plaque peptides were functionalized with TMT-2plex TMT-131 and footprint peptides were functionalized with TMT-2plex TMT-126. The peptide pools were then pooled and loaded onto a 100 µm internal diameter (ID) pre-column (Dionex) at 4 µL/min and separated over a 50 µm ID analytical column (C18 2 µm, Dionex). The peptides were eluted on a 0 to 35% acetonitrile gradient with an EASY-nLC 1000 nano-chromatography pump (Thermo Fisher, Odense Denmark). Data was acquired at 70,000 FWHM resolution in the MS mode and 17,500 FWHM resolution in the MS/MS mode. A total of 10 MS/MS scans were obtained per MS cycle.

### *In silico* methods for protein identification, selection, and sequence analysis

LC-MS/MS data was analyzed with PEAKS Studio 10.5 (Bioinformatics Solutions Inc.) and Scaffold 5.0 (Proteome Software Inc.). *De novo* sequences derived from MS/MS data were matched against the quagga mussel foot transcriptome (8) with tyrosine hydroxylation to DOPA, deamidation, oxidation, and carbamidomethylation set as variable modifications. Parent ion and fragment ion mass tolerances were set to 5 PPM and 0.01 Da, respectively. Acceptance criteria for identified proteins were peptide logP ≥ 15, protein –10 logP ≥ 50, and *de novo* average local confidence (ALC) score ≥ 80%, with at least two peptide spectra per identification.

Scaffold Q+ in Scaffold 5.0 was used to quantitate label-based quantitation peptide and protein identifications. Peptide identifications were accepted if they could be established at ≥ 75.0% probability to achieve an FDR < 0.1% and contained at least two identified peptides. Normalization against the reference (bulk plaque) were performed iteratively across samples and spectra on intensities, as described in (86). Spectra data were log-transformed and weighted by an adaptive intensity weighting algorithm.

Differentially expressed proteins were determined by applying the permutation test with unadjusted significance level p < 0.05 corrected by the Benjamini-Hochberg procedure.

The Protein Basic Local Alignment Search Tool (BLASTp) algorithm was applied on the identified proteins of interest against a clustered non-redundant database (38, 39) for homology assessment. Alignment of sequences were visualized in Jalview 2.11.3.3 using the Clustal Omega multiple sequence alignment program. The ProtParam online tool (Expasy) was used to calculate residue frequency, molecular weight, and isoelectric point of full-length protein sequences obtained from virtual translation of the open-reading frame of identified transcripts.

The method for analyzing tyrosine hydroxylation was categorized based on the level of confidence in the identified modification. A modification was considered confident if the tyrosine residue (Y) was highlighted with a specific residue number by the PEAKS PTM algorithm. It was classified as semi-confident if no number was indicated, but the spectrum contained a Y+16.99 m/z modification in both the b- and y-ions, indicating hydroxylation at that residue. Finally, a non-confident classification was assigned if a number was not indicated and the spectrum only showed the Y+16.99 m/z within an unfragmented peptide, suggesting that the tyrosine residue was not individually sequenced.

### Purification of Dbfp7

Dbfp7 was purified from quagga mussel phenol glands (**Figure S1**). The crude protein extraction method was adapted from (9). Phenol glands were dissected out of freshly harvested quagga mussels and homogenized in 5% acetic acid with 10 µM pepstatin and leupeptin protease inhibitors. The mixture was centrifuged at 20,000 x g for 40 minutes at 4°C. The pellet (P1) was then homogenized in 5% acetic acid and 8 M urea. The mixture is then centrifuged at 20,000 x g for 40 minutes at 4°C. The supernatant (S2), i.e. the urea-soluble proteins, was then mixed with mobile phase (0.1% trifluoroacetic acid (TFA)) and injected into a C4 reversed-phase column (XBridge Protein BEH C4, 300 Å). The eluent at 40 to 45% acetonitrile (with 0.1% TFA) was collected, lyophilized, and stored at -80°C until analysis.

### Structure Prediction with AlphaFold

The structure of Dbfp7 variant 1 was predicted with AlphaFold2 protein structure prediction (87). The query sequence was input in the ColabFold v1.5.5 python notebook and the resulting structures were visualized with ChimeraX v1.8.

### Gel electrophoresis

Samples were separated by molecular weight via gel electrophoresis in Bolt 12% Bis-Tris precast gels, using Bolt MES-SDS running buffer and Bolt LDS sample buffer. Gels were run at constant 165 V voltage for 40 minutes. The gels were fixed in 40% methanol + 10% acetic acid and stained with Coomassie Brilliant Blue R-250 for 1 hour. Gels were destained and visualized with a digital camera.

### ESI mass spectrometry

Dbfp7 was dissolved in 5% acetic acid at 400 µg/mL and ionized by electrospray ionization. The sample was analyzed with an Agilent 6538 UHD hybrid quadrupole time-of-flight mass spectrometer. The mass spectrum was processed using the Maximum Entropy algorithm implemented in the MassHunter BioConfirm software package.

### Nitroblue tetrazolium staining for DOPA detection

NBT was used to detect and quantify DOPA amounts. This protocol was adapted from (9, 88). Samples were prepared by adding 20 µL to 0.17 mM NBT in 2 M K- glycinate, then incubating at room temperature in the dark for 1 hour. The absorbance at 530 nm was measured and compared to a calibration curve generated using samples prepared with standard amounts of peptidyl DOPA. Peptidyl DOPA standards were synthesized by Biomatik and include the first 13 residues of Dbfp7, with the penultimate tyrosine replaced with DOPA.

Dot blots were prepared by depositing 10 µL of sample onto a nitrocellulose membrane and allowing it to dry in air for 30 minutes. The membrane was then rinsed with potassium glycinate for 30 minutes, then stained with 0.24 mM NBT for 1.5 hours in the dark, then rinsed with 0.2 M sodium borate for 1 hour.

### Sample preparation and spectra analysis for circular dichroism

To characterize the secondary structure of purified Dbfp7, far-UV circular dichroism was performed. The CD spectra was recorded using a Jasco spectropolarimeter, with a 2-nm bandwidth, scanning speed of 20 nm/min at a spectral resolution of 1.0 nm. The scans were averaged over ten scans, and the data was smoothed. The CD spectra of the buffer was recorded and manually subtracted from sample spectra. The secondary structure was characterized by computational analysis of the spectra using the Contin-LL program from the Dichroweb package (47).

Measured ellipticity was normalized to the molecular weight, pathlength, and concentration of the sample to get the mean residue ellipticity (deg*cm2/dmol). The average molecular weight of 7.2 kDa, as determined by mass spectrometry, was used.

### Sample preparation for atomic force microscopy

10 µL of sample was deposited onto freshly cleaved mica (V1 high grade) and incubated in a humidified chamber for 20 minutes. Residual protein was thoroughly washed away with blank solution (e.g. 5% acetic acid for Dbfp7 samples dissolved in 5% acetic acid). A droplet of deionized water was dropped onto the sample and placed in a fluid cell for AFM imaging in aqueous conditions.

### Imaging and adhesion force mapping via AFM PFQNM

Samples were imaged with a MultiMode Atomic Force Microscope equipped with a Nanoscope 6 controller operated via Nanoscope 10.0 software in PeakForce tapping mode with quantitative nanomechanical mapping (PFQNM). SCANASYST-FLUID probes with 20 nm (nominal) silicon nitride tips were calibrated on blank mica in water, resulting in calibrated spring constants around 0.7 N/m. The images were taken at 512- line resolution at 1.0 Hz line rate. ScanAsyst mode allows for automatic control of setpoint, gain, and scan rate. Initial images were taken with ScanAsyst auto-control on to determine the ideal setpoint for the sample. ScanAsyst auto-control was subsequently turned off to allow for direct pixel-to-pixel comparison of nanomechanical properties derived from the collected force-distance curves. PeakForce data was captured at a rate such that each height pixel has a corresponding force-distance curve saved. Images were analyzed using the Nanoscope Analysis 3.0 software. The individual curves for the entire map were baseline-corrected and moduli were fitted using the JKR contact model. Adhesion was calculated using the positive-slope algorithm.

## Author Contributions

A.R.O., R.H., D.J.R., J.N., and E.D.S. designed research; A.R.O., R.H., D.J.R., and J.N. performed research; A.R.O., R.H., D.J.R., J.N., K.M.M.C., and E.D.S. analyzed data; and A.R.O., R.H., K.M.M.C., and E.D.S. wrote the paper.

## Acknowledgements

This research was supported by Discovery Grant #342455 to E.D.S. from the Natural Sciences and Engineering Research Council of Canada (NSERC) (RGPIN-2019-06210), Post-Graduate Scholarship – Doctoral (PGS-D) to A.R.O. from NSERC, and the Queen Elizabeth II Graduate Scholarship in Science and Technology (QEII-GSST) to A.R.O. from the Government of Ontario. The authors thank the SickKids Proteomics, Analytics, Robotics & Chemical Biology Centre for assistance with LC-MS/MS data collection and interpretation, the Advanced Instrumentation for Molecular Structure Laboratory for ESI MS, and the SickKids Structural and Biophysical Core Facility for CD and HPLC. Special thanks are extended to Ryan Lee Chan for his assistance with AFM. The authors also gratefully acknowledge Dr. Magdalena Wojtas for her valuable insights and advice throughout the project.

## Abbreviations/Acronyms/Initialisms

DOPA: 3,4-dihydroxyphenylalanine
Dbfp: *Dreissena bugensis* foot protein
Dpfp: *Dreissena polymorpha* foot protein
MALDI-TOF: matrix-assisted laser desorption/ionization time-of-flight
LC-MS/MS: liquid chromatography tandem mass spectrometry
FP: footprint
BP: bulk plaque
IND: induced byssus
TMT: tandem mass tag
BLAST: basic local alignment search tool
KCl: potassium chloride
FCR: fold change ratio
HPLC: high-pressure liquid chromatography
NBT: nitroblue tetrazolium
pLDDT: predicted local distance difference test
AFM: atomic force microscope
CD: circular dichroism
BSA: bovine serum albumin
DMT: Derjaguin-Muller-Toporov
JKR: Johnson-Kendall-Roberts
SFA: surface forces apparatus
ID: internal diameter
TFA: trifluoroacetic acid

