## Supplemental Information for "Identification and characterization of a wet adhesive protein extracted from *Dreissena bugensis*, the freshwater quagga mussel"

**This PDF file includes:**

Supporting text
Figure S1
Figure S2
Figure S3
Figure S4
Figure S5
Figure S6
Table S1 title

**Other supporting materials for this manuscript include the following:**

Table S1 (full)

**Supporting Information Text**

### Calculation of Tabor Parameter

The Tabor parameter, $\mu_T$ is defined as follows [Tabor 1977; Kim 2024]:

$$\mu_T = \sqrt[3]{\frac{RW^2}{E^{*2}\varepsilon^3}}$$

Where

$$R = Radius\ of\ curvature$$
$$W = Work\ of\ adhesion$$
$$\varepsilon = intermolecular\ distance$$
$$\frac{1}{E^*} = \frac{1-\nu_1^2}{E_1} + \frac{1-\nu_2^2}{E_2}$$
$$E = Effective\ Young's\ modulus$$
$$\nu = Poisson's\ ratio$$

The intermolecular distance is assumed to be about 1.5 Å for carbon atoms at contact.

The radius of curvature of the SCANASYST-FLUID AFM probe tip is nominally 20 nm.

The Poisson's ratio for the two surfaces is assumed to be 0.25 and the effective Young's modulus for silicon nitride is 250 GPa and the effective Young's modulus for a thin protein film on mica is assumed to be about 1 GPa.

The work of adhesion is preliminarily determined to be about 6 x $10^{-4}$ N/m, the average work of adhesion calculated from the same adhesion force of 300 pN using DMT and JKR contact models of adhesion.

Altogether, $\mu_T$ is calculated to be 0.12.

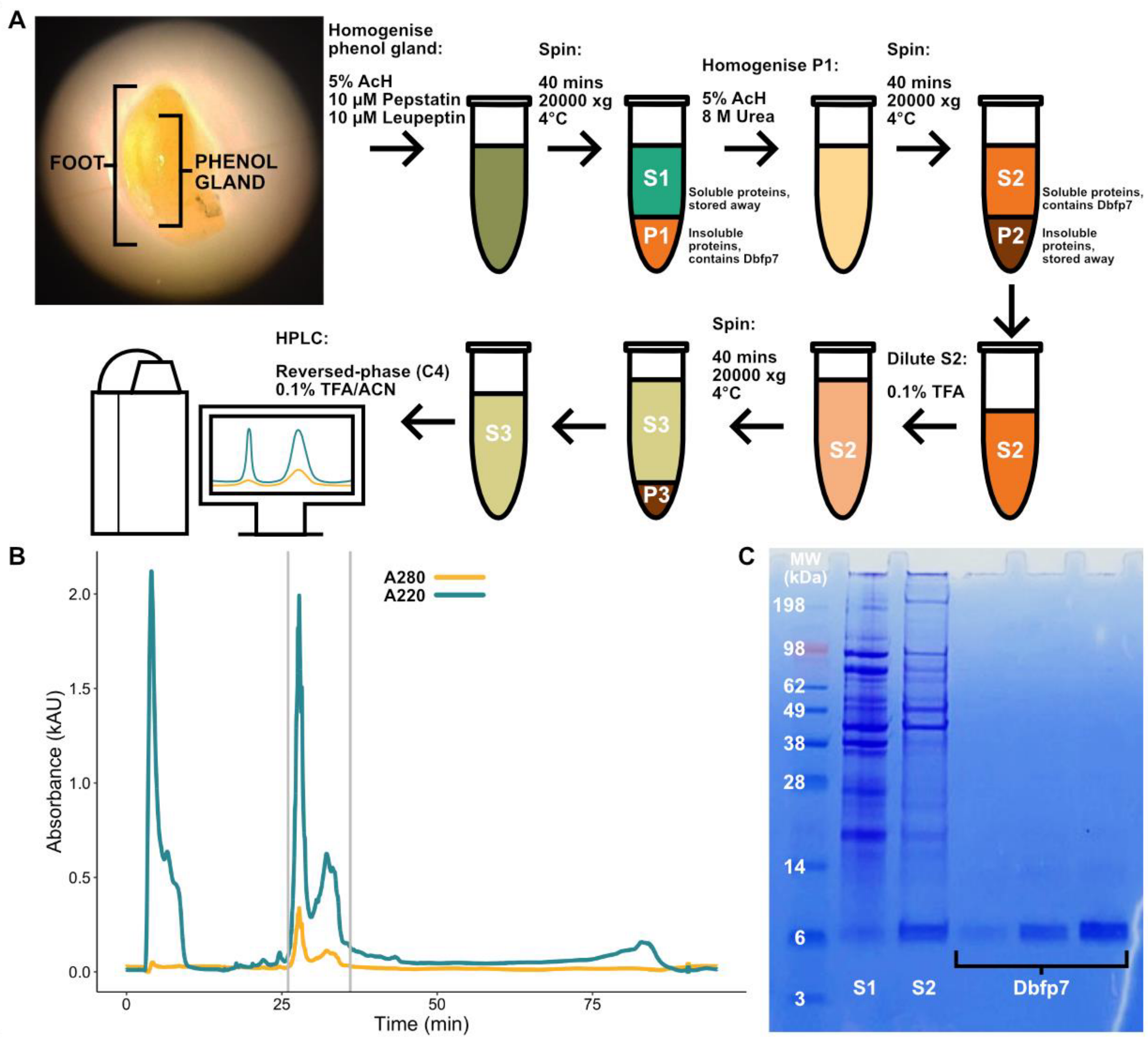


**Figure S1**: Extraction of Dbfp7. (A) Schematic of the extraction method. Phenol glands are dissected out of freshly harvested quagga mussels and homogenized in 5% acetic acid (Ach) with protease inhibitors, then subjected to several steps of selective precipitation, then purified with reversed-phase HPLC. (B) HPLC chromatogram of Dbfp7, within the vertical grey lines is the fraction containing Dbfp7. (C) Gel of phenol gland protein mixtures S1 and S2 and purified Dbfp7 increasing in concentration left to right.

**A**

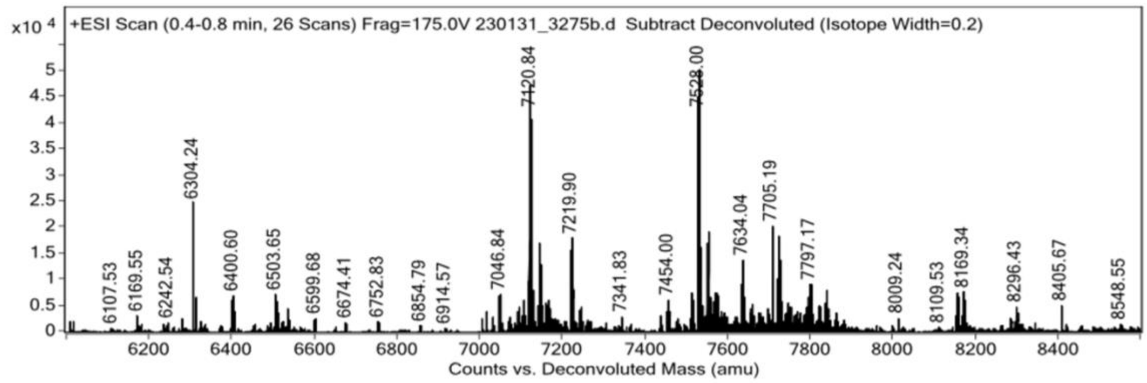


**B**

**Compound Mass List**

| # | Avg Mass | Abnd | MinZ | MaxZ | Z Count | Score |
|---|---|---|---|---|---|---|
| 1 | 7528.62 | 9 545.00 | 4 | 11 | 8 | 8.0 |
| 2 | 7121.08 | 9 434.00 | 3 | 9 | 7 | 9.0 |
| 3 | 7550.68 | 4 464.00 | 4 | 10 | 7 | 7.0 |
| 4 | 7721.51 | 4 262.00 | 4 | 11 | 8 | 6.0 |
| 5 | 7634.65 | 2 825.00 | 4 | 14 | 9 | 5.0 |

**C**

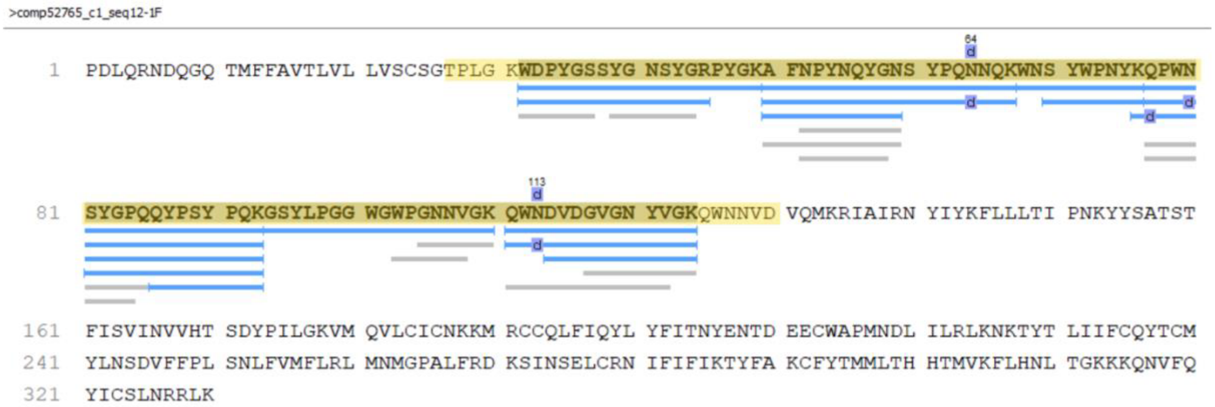


**Figure S1**: Mass spectrometric analysis of Dbfp7. (A) Deconvoluted MS spectrum of purified Dbfp7 with electrospray ionization. (B) Table of identified compound masses, indicating a majority of compounds are around 7 kDa in molecular weight. (C) Example coverage of Dbfp7 identified by LC-MS/MS. The entire virtually translated transcript is shown. Highlighted in yellow is the open reading frame of Dbfp7. Blue and grey lines indicate the peptides matched to the Dbfp7 sequence.

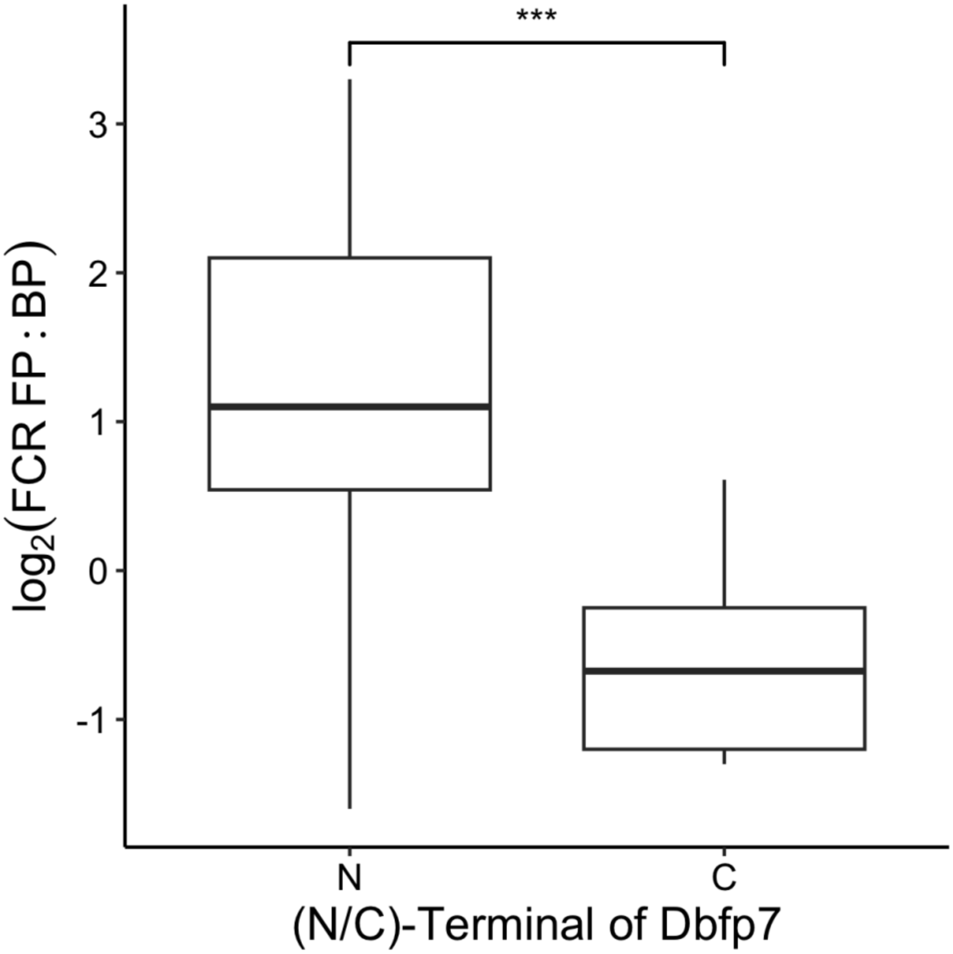


**Figure S2**: Relative enrichment of peptides of Dbfp7 detected by LC-MS/MS in either the N-terminal half or C-terminal half of the sequence. $\log_2$(FCR FP:BP) > 0 indicates peptides enriched in the footprint and $\log_2$(FCR FP:BP) < 0 indicates peptides enriched in the bulk plaque. *** significance p-value << 0.001

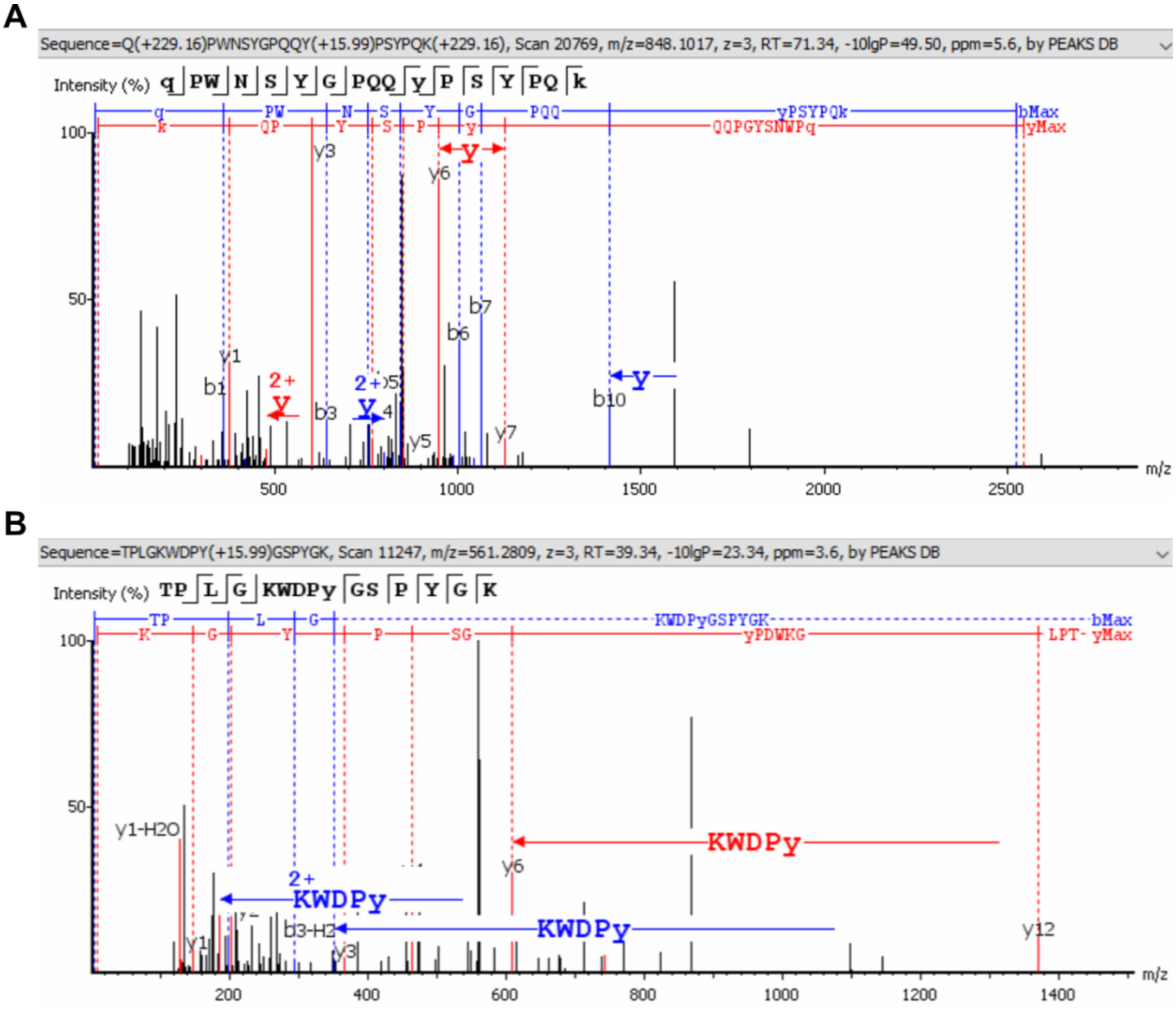


**Figure S4**: Example LC-MS/MS spectra of Dbfp7 peptides. (A) A peptide from variant 14 in the bulk plaque pool with a confident detection of DOPA (labelled lowercase ‘y’). (B) A peptide from variant 14 in the footprint pool with a non-confident detection of DOPA (labelled lowercase ‘y’ attached to the larger ‘KWDPy’ peptide).

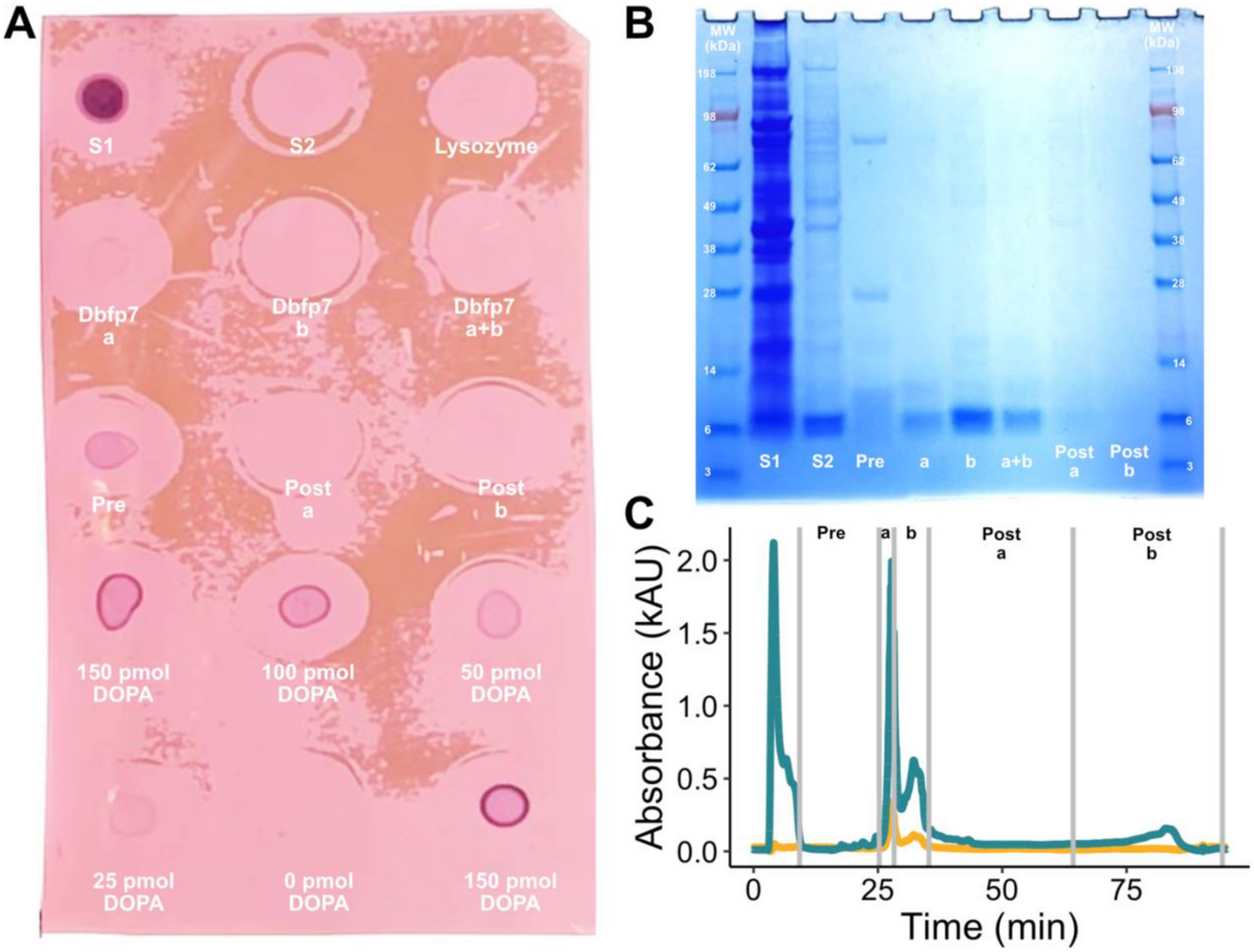


**Figure S5**: Dbfp7 is DOPA-deficient. (A) Nitroblue tetrazolium dot blot. S1 and S2 are protein mixture pre-HPLC. Lysozyme is the negative control. The second and third rows are HPLC fractions that were lyophilized then dissolved in 5% acetic acid, volume normalized by A220. The fourth and fifth rows are DOPA-containing peptides, with known amounts of DOPA loaded. (B and C) Corresponding gel and HPLC chromatogram.

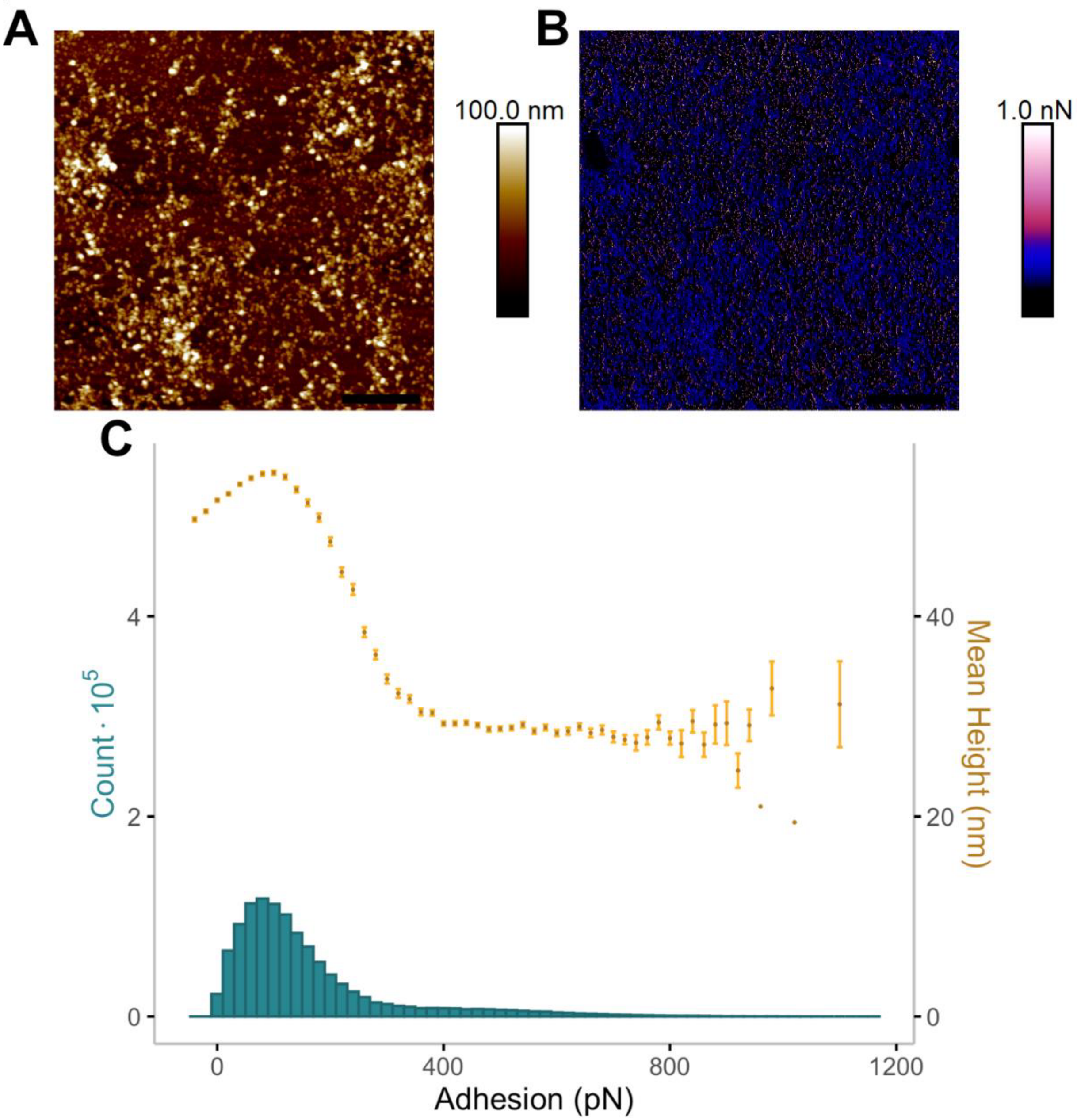


**Figure S6**: Atomic force microscopy of BSA at 100 µg/mL in water. (A) Height map. (B) Adhesion map. (C) The mean height of each bin in the adhesion histogram.

**Table S1**: Full list of identified proteins in induced byssus (IND) and naturally secreted bulk plaques (BP) and footprints (FP). Sector: the region of the identified protein, e.g. BPFP are proteins that are found in both bulk plaque and footprint. Accession #: transcript label and reading frame (0F, 1F, 2F, 0R, 1R, or 2R). Identification: protein identification determined by the BLASTp result. NR Clustered Database ID: top protein identified by BLASTp to an NR Clustered Database. E-value: estimate of the expected number of random alignments between the detected protein and the top BLASTp result. Other Significant Hits: other notable significant alignments to known proteins.

[Table attached as separate file entitled “Obille2024_TableS1.xlsx”]
